# LCM-seq identifies robust markers of vulnerable and resistant human midbrain dopamine neurons

**DOI:** 10.1101/2020.04.03.023770

**Authors:** Julio Aguila, Shangli Cheng, Nigel Kee, Ming Cao, Qiaolin Deng, Eva Hedlund

**Affiliations:** Department of Neuroscience, Karolinska Institutet, Stockholm, 171 77, Sweden; Department of Physiology and Pharmacology, Karolinska Institutet, Stockholm, 171 77, Sweden; Center for Molecular Medicine, Karolinska University Hospital, Stockholm, 171 77, Sweden

**Keywords:** Human midbrain dopamine neurons, spatial transcriptomics, laser microdissection, RNA sequencing, substantia nigra compacta, ventral tegmental area, Parkinson Disease

## Abstract

Defining transcriptional profiles of substantia nigra pars compacta (SNc) and ventral tegmental area (VTA) dopamine neurons in human is critical to understanding their differential vulnerability in Parkinson Disease. However, reported marker profiles for these neuron populations are derived predominantly from rodents, utilize small sample sizes and display extensive variability between studies. Here, we map selective expression profiles of dopamine neurons in an extensive collection of human SNc and VTA using laser capture microdissection coupled with Smart-seq2 RNA sequencing (LCM-seq). By applying a bootstrapping strategy as sample input to DESeq2, we identify 33 differentially expressed SNc- or VTA-specific markers and we also compute the minimal cohort size required to identify differentially expressed genes (DEGs) that are concordant regardless of cohort size. Among the identified DEGs, *ZCCHC12, CDH13* and *SERPINE2*, are minimally required to distinguish SNc or VTA dopamine neurons in both human and mouse. In summary, our study identifies novel markers, besides previously identified ones, which will be instrumentsal for future studies aiming to modulate dopamine neuron resilience as well as validate cell identity of stem cell-derived dopamine neurons.

## INTRODUCTION

Midbrain dopamine neurons are divided into two major populations, the substantia nigra pars compacta (SNc) and the ventral tegmental area (VTA) (Hedlund and Perlmann 2009). SNc dopamine neurons project to the dorsolateral striatum (Dahlstroem and Fuxe 1964) and are severely affected in Parkinson Disease (PD) (Damier et al. 1999b; Damier et al. 1999a), while VTA dopamine neurons project to cortical and mesolimbic areas and are more resilient to degeneration (Hedlund and Perlmann 2009). These neuron populations have been extensively investigated in numerous rodent models, (Grimm et al. 2004; Chung et al. 2005; Greene et al. 2005; Bifsha et al. 2014; Poulin et al. 2014), towards the goal of identifying molecular mechanisms that can prevent degeneration or to model disease. Targeted analysis of midbrain dopamine neuron populations have revealed several markers that appear to differentially label SNc e.g. *Aldh1a7, Sox6, Cbln1, Vav3, Atp2a3* and VTA e.g. *Calb1, Otx2, Crym, Cadm1* and *Marcks* (Damier et al. 1999a; Grimm et al. 2004; Chung et al. 2005; Greene et al. 2005; Di Salvio et al. 2010; Bifsha et al. 2014; Panman et al. 2014; Nichterwitz et al. 2016). Transcriptional analysis of human tissue has largely been limited to SNc (Cantuti-Castelvetri et al. 2007; Simunovic et al. 2009) except for a recent small sample cohort of both SNc and VTA (Nichterwitz et al. 2016). Frustratingly, these aforementioned investigations display extensive cross-study variability, resulting in very few reproducible markers both within the same species and across mouse, rat and human. Small sample sizes could be confounding these findings, along with differences in rodent strain backgrounds, inter-individual variability among human patients, and methodological differences. Thus, these discrepancies warrant further comprehensive cross-species comparative analyses.

To identify human SNc and VTA specific markers, which could reveal cell intrinsic properties underlying their differential vulnerability in PD, a thorough large-scale transcriptional profiling in adult human tissues is required. Such an analysis could also investigate the minimum cohort size necessary, above which lineage specific markers remain stably differentially expressed irrespective of patient selection, an essential requirement for valid study design in variable human populations. Finally, identified differences could also serve as a foundation for the selective *in vitro* derivation of SNc dopamine neurons, which represent the ideal cell type for transplantation in PD (Schultzberg et al. 1984; Haque et al. 1997; Thompson et al. 2005; Hedlund and Perlmann 2009; Kriks et al. 2011; Ganat et al. 2012).

Here we used the spatial transcriptomics method LCM-seq, which combines laser capture microdissection with Smart-seq2 RNA sequencing (Nichterwitz et al. 2016; Nichterwitz et al. 2018), to precisely analyze individually isolated SNc and VTA dopamine neurons from 18 human *post mortem* brains. Using a bootstrapping without replacement coupled with DESeq2, we identify 33 markers that were stably differentially expressed between SNc and VTA dopamine neurons. We show that the minimal cohort size required to reliably identify these subtype-specific markers were eight subjects, which may explain why smaller cohorts have given inconsistent results. We further applied this approach to a large-scale data set of mouse single midbrain dopamine neurons, revealing 89 SNc and VTA mouse stable genes. Finally, we show that only three of the human stable genes, *ZCCHC12, CDH13* and *SERPINE2*, overlap with the mouse stable genes, but that these three markers nevertheless faithfully classify SNc or VTA dopamine neurons in both species. Several of the markers identified here have been implicated in PD or other degenerative diseases and thus provide compelling future targets to modulate neuronal vulnerability or to model disease.

## RESULTS

### Published inter- and intra-species SNc and VTA transcriptional profiles display considerable discrepancies

To begin constructing a consensus for VTA and SNc specific molecular profiles, we compiled previously published transcriptome data from mouse and rat, for VTA or SNc differentially expressed genes (DEGs) (Grimm et al. 2004; Chung et al. 2005; Greene et al. 2005; Bifsha et al. 2014; La Manno et al. 2016). This analysis revealed that a surprisingly low fraction of DEGs were common across data sets (Supplemental Fig. S1A, B; Supplemental Tables S1 and S4). Comparing across species with our previously published small data set on human SNc and VTA (Nichterwitz et al. 2016), only two genes, *SOX6* and *CALB1*, overlapped within SNc and VTA gene lists, respectively (Supplemental Fig. S1C). These discrepancies highlighted the urgent need to identify reproducible marker profiles for VTA and SNc dopamine neurons.

### LCM-seq derives SNc or VTA specific markers from a large human cohort

To identify robust and specific human dopamine neuron subpopulation markers, we isolated individual VTA and SNc neurons from *post mortem* tissues of 18 adult individuals by LCM (Supplemental Fig. S2A-G; Supplemental Table S2) and conducted polyA-based RNA sequencing. This study represents the largest human data set profiling of SNc and VTA dopamine neurons to date. The quality of human fresh frozen tissues used may vary as a consequence of *post mortem* interval (PMI), sample handling and preservation. Therefore, prior to conducting differential gene expression analysis we performed extensive quality control analysis to rule out undesired influences from sample processing (Supplemental Table S5). Randomly selected samples that exhibited different PMIs for VTA or SNc neurons displayed comparable cDNA quality (Supplemental Fig. S2H, I). Furthermore, while the total number of reads varied between individual samples, such variability was similarly distributed between SNc and VTA samples (Supplemental Fig. S2J). The number of detected genes did not correlate with either the age of the donor (Supplemental Fig. S2L), the PMI (Supplemental Fig. S2M) or the total number of reads (Supplemental Fig. S2K). Only the number of collected cells per sample modestly impacted gene detection (P=0.515) (Supplemental Fig. S2N). However, neither the number of collected cells nor the number of detected genes were significantly different between SNc and VTA neuron groups (Supplemental Fig. S2O, P) and thus should not affect DEG identification. Finally, we observed that all samples strongly expressed the dopamine neuron markers *EN1/2, FOXA2, LMX1B, PITX3, NR4A2, TH* and *SLC6A3* (DAT), and the general neuronal marker NEFH, while lacking glial markers *MFGE8, CX3CR1* or *GPR17*. This clearly demonstrates the selective enrichment of dopamine neurons using the LCM-seq methodology (Fig. 1A). *KCNJ6* (GIRK2) and *CALB1*, two genes often used to distinguish between SNc or VTA dopamine neurons, were also expressed (Fig. 1A), but interestingly could not, on their own, accurately classify our samples (Supplemental Fig. S2Q).

**Figure 1.**
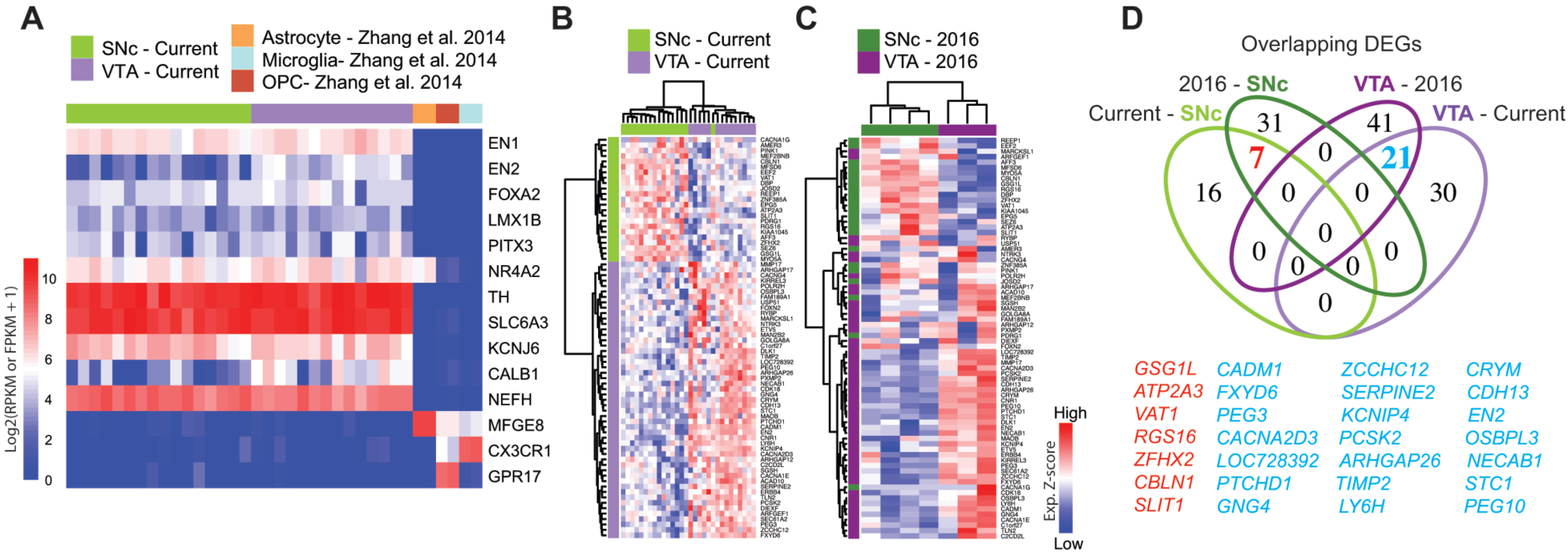
Gene signature of human adult midbrain dopamine neurons using LCM-seq. (A) The high sample quality for the 18 male subjects profiled in this study was confirmed by strong expression of the midbrain dopamine neuron markers *EN1/2, FOXA2, LMX1B, PITX3, NR4A2, TH* and *SLC6A3* (DAT), the pan-neuronal marker neurofilament (*NEFH*), and the lack of astrocyte, microglia or oligodendrocyte precursor contamination (Zhang et al. 2014). (B) Hierarchical clustering analysis of samples from the current study using the 74 DEGs identified by DESeq2. (C) The 74 DEGs also separated SNc and VTA samples from Nichterwitz et al., 2016 (3 female subjects). (D) Venn-diagram displaying an unexpected low degree of overlap between SNc and VTA DEGs found independently in each study. See also Supplemental Fig. S2.

Differential expression analysis, considering these 18 individuals, identified 74 DEGs (Supplemental Table S6), which resolved SNc from VTA neurons (Fig. 1B). These genes also distinguished SNc from VTA samples from a small cohort (N=3) investigated previously (Nichterwitz et al. 2016) (Fig. 1C). However, few overlapping DEGs were identified between these two studies, even though the same experimental method was used. In fact, only seven and 21 DEGs among the SNc and VTA gene lists, respectively overlapped (Fig. 1D). Notably, the 100 DEGs identified in the small cohort (Nichterwitz et al. 2016) (Supplemental Fig. S2R), failed to distinguish SNc and VTA for the current larger cohort of 18 subjects (Supplemental Fig. S2S; Supplemental Table S7). This suggests that sample size affects identification of DEGs and that a larger cohort size may identify DEGs that more robustly distinguishes the two populations. Moreover, this straight-forward DESeq2 analysis, that considers all samples together, cannot address how significant the 74 DEGs would be when considering smaller patient subsets from the cohort.

### Bootstrapping coupled with DESeq2 identifies stable DEGs unique to human SNc or VTA

To evaluate how cohort subsets may affect DEG detection, we used a bootstrapping algorithm in combination with DESeq2. Importantly, to reduce the biological variability we considered only those subjects for which both SNc and VTA samples were available (12/18 subjects, 24 samples in total). To begin with this approach, a subset of three subjects were randomly chosen from the pool of total 12 subjects. Differential expression analysis was then performed between the SNc and VTA samples of these subjects (DESeq2), and genes were scored as differentially expressed (DE) or not (adj P-value < 0.05). This random sampling of three subjects, followed by DESeq2 analysis, was performed a total of 1000 times, and the DE frequency over these 1000 comparisons was recorded for this iteration (i = 3 subjects). Subsequently, this process was repeated using subsets of four subjects, then five, up to a maximum of 11 of the 12 subjects.

For each subset size (i^3^ to i^11^) the DE frequency was recorded for each gene over the 1000x comparisons of that iteration (Fig. 2A). By considering all DE genes in an iteration (1000x comparisons), we were able to detect hundreds of genes, much more than were detected in any individual comparison, where on average only e.g. 13 DE genes were detected in i^3^ (Supplemental Fig. S3A). Interestingly, we found that eight subjects were required to saturate the sensitivity of our gene detection, as few new genes were identified when considering additional subjects in subsequent iterations (Fig. 2B).

**Figure 2.**
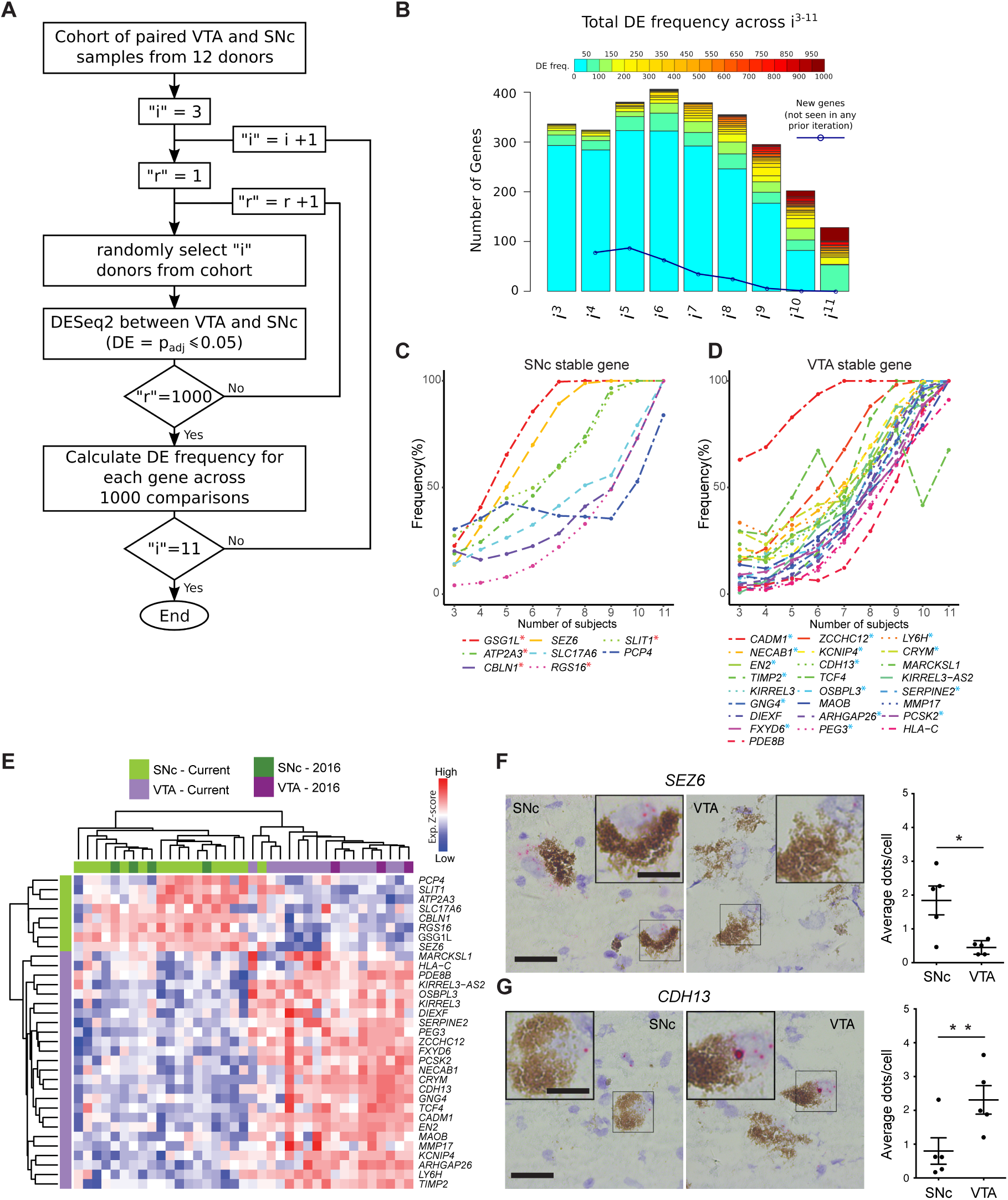
Bootstrapping analysis coupled with DESeq2 identifies SNc and VTA stable genes from a cohort of 12 human subjects. (A) Schematic of the analysis workflow. (B) Numbers of genes displaying a given DE frequency in each iteration of the workflow, from i = 3 subjects, up to i = 11 subjects. (e.g. in i^3^, almost 300 genes were DE between 1-50 times out of 1000 x r). (C, D) Stable gene DE frequency increased at each iterative sample size increase, for SNc (C) or VTA (D). For each gene list, genes that overlap with DEGs from Figure 1D are marked with an asterisk (*). (E) Hierarchical clustering analysis using the 33 stable genes, faithfully segregated both SNc and VTA samples. (F, G) RNAscope staining and quantification for the stable genes *SEZ6* (F, p=0.025) and *CDH13* (G, p=0.005) enriched in the SNc and VTA, respectively (n=5 subjects, data represented as mean ± SEM, Paired t test). Scale bars 30 μm (15 μm for insets). See also Supplemental Fig. S3.

We then summed the DE frequency across i^3^ to i^11^ (9000 comparisons in total), separated genes into SNc or VTA enriched lists, and ranked the lists from most to least frequently DEs (Supplemental Fig. S3C, D; Supplemental Table S8). Highly ranked genes on these two lists included multiple known human (e.g. *GSG1L, SLIT1, ATP2A3, CADM1, CRYM, TCF4*) and rodent rodent (e.g. *Gsg1l, Slit2, Atp2a3, Cadm1, Tcf4*) SNc and VTA markers (Grimm et al. 2004; Chung et al. 2005; Greene et al. 2005; Cantuti-Castelvetri et al. 2007; Simunovic et al. 2009; Bifsha et al. 2014; Poulin et al. 2014; La Manno et al. 2016; Nichterwitz et al. 2016). To identify the most reliable DE genes, we designated genes that were detected more than 3000 times (out of 9000) as “stable genes”. This stringent cutoff was chosen since the resulting SNc and VTA lists would then contain at least one stable gene that could be identified during the first iteration (where three individuals were used as the sample size).

We identified eight stable genes for the SNc (Fig. 2C) and 25 stable genes for the VTA (Fig. 2D). Five SNc (labeled in red**\***: *GSG1L, ATP2A3, CBLN1, RGS16* and *SLIT1*) and 15 VTA (in blue**\***: *CADM1, NECAB1, EN2, TIMP2, GNG4, FXYD6, ZCCHC12, KCNIP4, CDH13, OSBPL3, ARHGAP26, PEG3, LYH6, CRYM, SERPINE2* and *PCSK2*) stable genes were among the aforementioned 7 and 21 overlapping DEGs, identified across the two studies (Fig. 1D). We also compared this stable gene list with the outcome of DESeq2 analysis alone, when applied to the same 12 subjects (Supplemental Fig. S3E, F). Reassuringly, all 8 SNc and 25 VTA markers perfectly overlapped with the DEGs from DESeq2 alone using an adjusted *P*-value <0.05 (Supplemental Fig. S3E) or a more stringent significance (adj. *P*-value <0.01, Supplemental Fig. S3F). The expression of SNc stable genes was confirmed in two independent human microarray datasets (which lacked VTA samples) (Supplemental Fig. S3G) (Cantuti-Castelvetri et al. 2007; Simunovic et al. 2009). Further, we used RNAscope (Supplemental Fig. S3H) (Wang et al. 2012) to validate the expression pattern of the SNc stable gene *SEZ6*, and the VTA stable gene *CDH13*, in *post mortem* human tissues (Fig. 2F-G), further ratifying our LCM-seq data and the bootstrapping approach. Importantly, the stable genes faithfully classified SNc and VTA from 21 individuals (Fig. 2E), namely all 18 male individuals from our current dataset and the three female samples investigated previously (Nichterwitz et al. 2016).

In conclusion, we have identified 33 markers that correctly classify samples as either SNc or VTA, and that remain robust to individual subject variability. Notably, these genes were stably differentially expressed only when at least eight subjects were included in the bootstrapping strategy (Fig. 2D, E), thereby defining a minimal cohort size required to distinguish SNc and VTA samples in human subjects using LCM-seq.

### Common stable genes can classify SNc and VTA dopamine neurons across species

To identify common stable DEGs for VTA and SNc across other species, we applied the combined bootstrapping and DESeq2 strategy to a single-cell RNA-sequencing (scRNA-seq) dataset that profiled postnatal mouse midbrain dopamine neurons (Supplemental Fig. S4A, raw data analyzed here) (La Manno et al. 2016). Single cells were initially quality controlled for expression of known dopamine neuron markers, and the absence of contaminating glia or oligodendrocytes markers (Supplemental Fig. S4B) (Zhang et al. 2014), followed by iterative bootstrapping and DESeq2 analysis (Supplemental Fig. S4C).

This identified 36 SNc- and 53 VTA-specific mouse stable genes (Fig. 3A, B; Supplemental Table S9, Supplemental Fig. S4D, E, red labels) including novel genes (e.g. *Pde2a, Adrbk2, Tppp3, Col25a1, Rph3a, Mgst3*) in addition to previously reported markers (e.g. *Atp2a2, Cadm2, Serpine2, Zcchc12, Ly6e, Calb1*) (Grimm et al. 2004; Chung et al. 2005; Greene et al. 2005; Bifsha et al. 2014; Poulin et al. 2014; La Manno et al. 2016; Nichterwitz et al. 2016). Multiple known SNc and VTA markers ranked just below our 30% frequency cutoff and were thus not designated as stable genes (Supplemental Fig. S4D, E, black labels). Importantly, the 89 stable genes correctly classified the mouse single cells as either SNc or VTA (Supplemental Fig. S4F).

**Figure 3.**
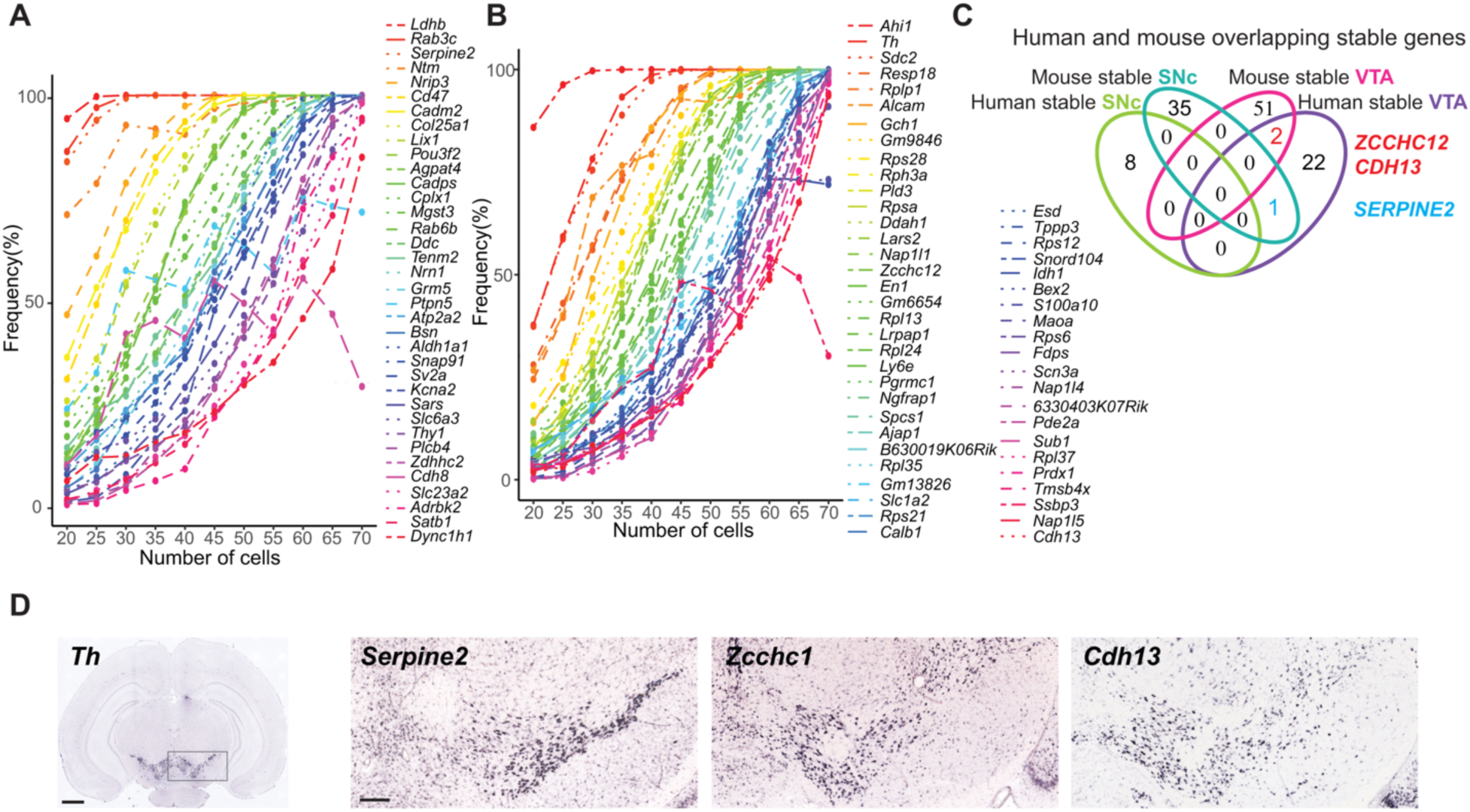
Bootstrapping analysis coupled with DESeq2 identifies SNc or VTA stable genes from a mouse scRNA-seq dataset. (A, B) Identification of SNc (A) and VTA (B) stable genes from 146 single mouse dopamine neurons (La Manno et. al, 2016) (73 available SNc cells and 73 randomly selected VTA cells), see METHODS. (C) Venn-diagram displaying overlapping stable genes between mouse and man. The three common markers *ZCCHC12, CDH13* and *SERPINE2* are highlighted in red. (D) *In situ* hybridization images of coronal midbrain sections of the adult mouse (P56, Allen Brain Atlas) showing RNA expression pattern for *Th* and the common stable genes *Serpine2, Zcch12* and *Cdh13*. Scale bars: (D) 1mm for *Th* and 200 μm for remaining markers. See also Supplemental Fig. S4.

We next cross compared our human and mouse stable gene lists. Two genes, *ZCCHC12* and *CDH13*, were common to human and mouse VTA. In addition, *SERPINE2* was expressed in human VTA, while it’s mouse homolog *Serpine2*, was expressed in the mouse SNc (Fig. 3C). The expression patterns of these three genes were corroborated in the adult mouse using Allen *in situ* images (Fig. 3D). To investigate the predictive power of these three common genes, we computationally classified human LCM-seq or mouse scRNA-seq samples using the human and mouse stable gene lists, while including or omitting the three common genes (Fig. 4A-H). (100% reflects perfect classification accuracy, while 50% reflects a random outcome).

**Figure 4.**
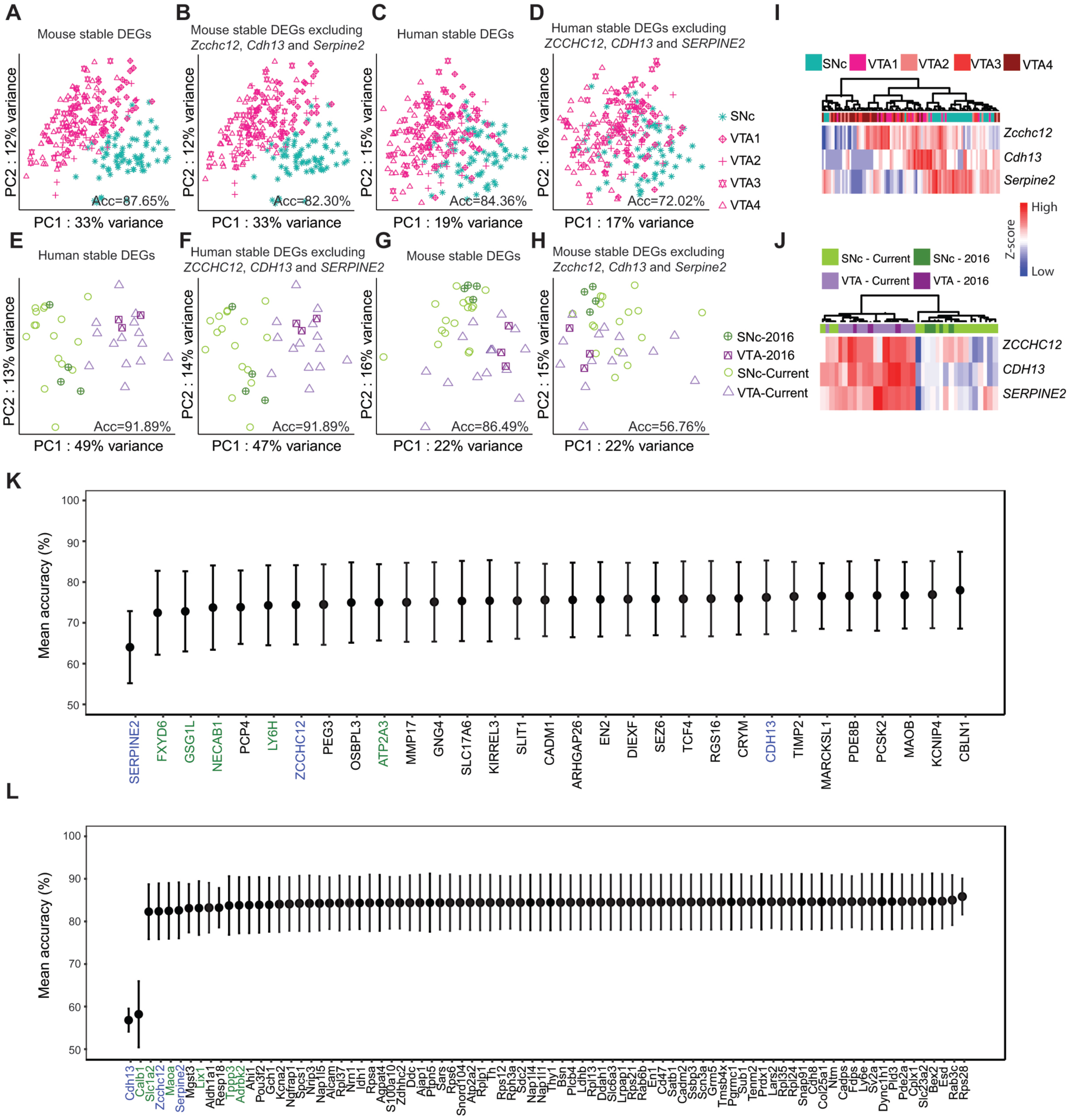
Predictive power of mouse and human stable genes to resolve SNc and VTA across species. (A-D) PCAs of the mouse single cells when considering: all mouse stable genes (A), mouse stable genes excluding the three common genes (B), human stable genes (C) or human stable list excluding the three common genes (D). (Classification accuracy = Acc%) (E-H) Similar analysis as A-D but applied to 21 human subjects (includes Nichterwitz et al., 2016, subjects). (I, J) Hierarchical clustering analysis considering only the three common stable genes are sufficient to classify SNc and VTA samples in mouse (I) and human (J). (K) Ranked mean accuracy for individual human stable genes to resolve the mouse dataset, upon random exclusion together with any other two stable genes. The three common stable genes between mouse and human are labelled in blue, while additional genes with predictive value in the mouse are labelled in green. (L) Ranked mean accuracy for individual mouse stable genes to resolve the human dataset, upon random exclusion together with any other two stable genes. Common stable genes between mouse and human are labelled in blue while additional genes with some predictive value in the human (led by *Calb1*) are labelled in green. See also Supplemental Fig. S5.

When using a complete stable gene list to resolve samples within the same species, negligible reduction in the accuracy was observed after removal of the three common genes in either mouse (Fig. 4A, B) or human (Fig. 4E, F). Despite the low overlap of stable genes across mouse and human, they could to some extent separate samples from the other species (Fig. 4C, G). However, subsequent exclusion of the three common genes led to a reduction in classification accuracy (Fig. 4D, H). This was most pronounced when attempting to classify human samples using the mouse stable genes excluding *Zcchc12, Cdh13* and *Serpine2*, resulting in an almost chance accuracy of 56.76% (Fig. 4H). Reassuringly, these three markers alone were able to largely segregate SNc from VTA samples in both species (Fig. 4I, J). In conclusion, we identified 89 markers for the mouse SNc and VTA using the combined bootstrapping and DESeq2 approach. Cross-species analysis further revealed three mouse stable genes overlapping with the human stable gene list, that alone are sufficient to correctly classify these two neuron subpopulations in both species.

### Systematic analysis of classification accuracy identifies additional predictive SNc and VTA markers

We next sought to systematically interrogate if the remaining stable human and mouse genes that, despite not overlapping between species, could prove useful in cross-species classification. For a given species, we randomly excluded any three stable genes and calculated the mean accuracy of the remaining markers in resolving the other species’ SNc and VTA samples (Fig. 4K, L). Our random exclusion paradigm demonstrated that *SERPINE2* was the most important gene resolving mouse SNc and VTA samples, as it occupied the top rank, while *ZCCHC12* and *CDH13* occupied ranks 7 and 24, respectively (Fig. 4K). Interestingly, the human stable genes *FXYD6, GSG1L, ATP2A3, NECAB1* and *LY6H*, were also highly ranked in their accuracy to classify mouse samples (Fig. 4K). The expression patterns of these genes were corroborated in the adult mouse using Allen *in situ* images (Supplemental Fig. 3D and S5A).

Unsurprisingly, *Cdh13, Zcchc12* and *Serpine2* were top ranked (1, 4 and 6 respectively) upon exclusion from the mouse stable gene list when resolving human samples (Fig. 4L). However, only *Cdh13* strongly contributed to correct sample classification, along with *Calb1*, a known VTA marker in both mouse (Chung et al. 2005; Greene et al. 2005; La Manno et al. 2016) and human (Reyes et al. 2012; Nichterwitz et al. 2016). Accordingly, the rank of *CALB1* on the human VTA list was close to the 30% frequency threshold for the “stable gene” classification (Supplemental Fig. S3B; Supplemental Table S8).

The exclusion of other mouse stable genes, including *Slc1a2, Maoa, Tppp3, Lix1* and *Adrbk2*, which also fell outside of the 30% frequency threshold for human stable genes (Supplemental Table S8), barely impaired the classification of human samples (Fig. 4L). Similar to *Calb1, in situ* analysis of these genes in the adult mouse was consistent with their differential expression between VTA and SNc (Supplemental Fig. S5B, green labels). Many novel mouse stable genes, while displaying the expected expression patterns in line with their SNc or VTA enrichment (Supplemental Fig. S5B, black labels), had no impact on the classification of human samples (Fig. 4L). In conclusion, we identified additional human stable genes, including *FXYD6, GSG1L, ATP2A3, NECAB1* and *LY6H* with power to also distinguish SNc or VTA dopamine neurons in mouse.

## DISCUSSION

The selective vulnerability of SNc dopamine neurons to PD, and the relative resilience of VTA dopamine neurons, has encouraged the field to investigate the molecular signature of these two neuron subpopulations. When we analyzed existing data sets (Grimm et al. 2004; Chung et al. 2005; Greene et al. 2005; Bifsha et al. 2014; Poulin et al. 2014; La Manno et al. 2016; Nichterwitz et al. 2016), we identified large discrepancies in the reported SNc or VTA enriched genes across different studies. Such inconsistencies could result from multiple factors, including small sample sizes, as well as variability between subjects, which is recognized to be a major confounding factor in human studies (Mele et al. 2015). This encouraged us to conduct a large focused study on adult human midbrain dopamine neurons using LCM-seq (Nichterwitz et al. 2018). We consequently constructed a comprehensive LCM-seq dataset, isolating single SNc or VTA dopamine neurons from *post mortem* tissues of total 18 individuals, the largest collection of human dopamine neurons, aiming to reveal robust molecular signatures to distinguish the two subpopulations.

Using an iterative bootstrapping without replacement coupled with DESeq2 (available at https://github.com/shanglicheng/BootstrappingWithoutReplacement), and a strict selection criteria (here, a 30% threshold for stable classification) we identify 33 of the most stable DEGs. 25 of these genes define VTA identity, while eight define SNc identity, and together they can accurately classify LCM-seq samples from our previous (3 females), and current (18 males) patient cohorts. Importantly, we also find that a minimum of eight subjects are required to identify these human stable genes using our criteria, and we speculate that identifying lineage-specific markers between any two highly related cellular sub-populations will also require carefully considered, and appropriately large cohort sizes. Such considerations also apply to studies comparing, for example, healthy and diseased dopamine neurons, that may exhibit potentially subtle pathological changes.

Application of this approach to a mouse single-cell dataset profiling midbrain dopamine neurons (La Manno et al. 2016), identified 89 stable DEGs, among which *Zcchc12, Cdh13* and *Serpine2* overlapped with the human stable gene list. Unexpectedly, these genes alone were sufficient to classify VTA and SNc dopamine neurons in both species, which also validated our approach to enrich robust DEGs. These three genes were so far unexplored in the dopamine system. Serpine2 (Glia-derived nexin) is a serine protease inhibitor which can promote neurite extension by inhibiting thrombin, and which appears downregulated in Alzheimer’s disease (Choi et al. 1995). Serpine2 promotes biogenesis of secretory granule which is required for neuropeptide sorting, processing and secretion (Kim and Loh 2006). *Cdh13* encodes for adhesion protein 13, which together with other family members as *Cdh9* and *Cdh15* are linked to neuropsychiatric disorders (Redies et al. 2012). Cdh13 can regulate neuronal migration and also has an effect on axonal outgrowth as demonstrated in the serotonergic system (Forero et al. 2017). Zcchc12 on the other hand is a transcriptional co-activator that positively regulates BMP signaling through interaction with Smads and CBP, and appears necessary for basal forebrain cholinergic neuron gene expression (Cho et al. 2008b). When mutated, Zcchc12 appears to cause X-linked mental retardation (Cho et al. 2008a). It remains to be investigated how dopamine neurons respond to the loss of any of these three factors.

Remarkably, the species-specific gene lists could also partly classify SNc and VTA dopamine neurons of the other species. Moreover, some stable genes’ closely related family members are also expressed in SNc and VTA samples, e.g. *Atp2a2, Cadm2, Ly6e, En1, Maoa* and *Pde2a* are present in the mouse while *ATP2A3, CADM1, LY6H, EN2, MAOB* and *PDE8B* are found in human. Further, our analysis also demonstrates important differences between the human or mouse stable genes. *Serpine2* was expressed in mouse SNc, while *SERPINE2* was expressed in human VTA. In addition to the three common markers, several human stable genes (e.g. *FXYD6, GSG1L, ATP2A3, NECAB1 and LY6H*) contribute in classifying mouse SNc and VTA samples. Hence, there remains a solid rational to continue investigating these human-relevant genes in the mouse.

The stable DEGs we identified here may be highly relevant to induce resistance or model disease as previously attempted in rodents (Chung et al. 2005; Poulin et al. 2014). Several of the human stable genes (or related family members) e.g. *GSG1L, ATP2A3, SLC17A6, RGS16, KCNIP1, CDH13, TCF12, OSBPL1A, OSBPL10, GNG7, ARHGAP18, ARHGAP24, PCSK5, PEG3, HLA-DOA, HLA-DRA, HLA-DRB1* and *PDE8B* have been found dysregulated in PD (Cantuti-Castelvetri et al. 2007; Bossers et al. 2009; Simunovic et al. 2009) and/or are represented in PD datasets from genome wide association studies (GWASdb SNP-Disease Associations dataset, http://amp.pharm.mssm.edu). Interestingly, mice lacking *Rgs6*, a related family member of the human SNc stable gene *RGS16*, develop specific degeneration and cell loss of SNc dopamine neurons at the age of 12 months (Bifsha et al. 2014). Loss of the SNc stable gene *Cplx1* results in a compromised nigrostriatal pathway in knockout mice (Hook et al. 2018). Moreover, mutations in the human SNc stable gene *SEZ6* have been implicated in diseases such as Alzheimer’s (Khoonsari et al. 2016; Paracchini et al. 2018), childhood-onset schizophrenia (Ambalavanan et al. 2016), epilepsy and febrile seizures (Yu et al. 2007; Mulley et al. 2011).

Regarding cell replacement therapies targeting PD (Kriks et al. 2011; Ganat et al. 2012; Hallett et al. 2014; Kefalopoulou et al. 2014; Kirkeby et al. 2017), there remains an urgent need to optimize the pluripotent stem cell preparations to specifically generate SNc rather than VTA neurons (Barker et al. 2017; Sonntag et al. 2018). Evaluation of the correct patterning and differentiation of pluripotent cells to midbrain dopamine neurons relies upon gene expression analysis using quantitative real time PCR (qPCR) or global transcriptome approaches such as RNA sequencing (Ganat et al. 2012; Barker et al. 2017; Nolbrant et al. 2017; Studer 2017). Hence, accurate reference gene signatures of adult human SNc neurons are critical towards further advancements in the regenerative PD field. Our LCM-seq and computational stable gene analysis can therefore serve as a base criterion describing the transcriptional profile of adult, human SNc and VTA neurons. This will greatly facilitate dopamine neuron replacement efforts, in addition to disease modeling studies using dopamine neurons derived from patient-specific pluripotent cells (Miller et al. 2013; Vera et al. 2016).

In summary, using LCM-seq and a bootstrapping approach coupled with DESeq2, we have identified SNc and VTA dopamine neuron markers in both human and mouse. We reveal that the smallest human cohort required to detect such stable DEGs consists of eight individuals, informing future study designs targeting highly related cellular populations. We find that only three genes are equally and strongly predictive across the two species, highlighting the need to carefully define human dopamine neuron subpopulations even when mouse marker profiles are available. This human transcriptomic data set, derived from individually isolated dopamine neurons, will thus help further our understanding and modeling of selective neuronal vulnerability and resilience, and serve as a reference for derivation of authentic SNc or VTA dopamine neurons from stem cells.

## METHODS

### Ethics statement

We have ethical approval to work with human *post mortem* samples (Supplemental Tables S2 and S3) from the regional ethical review board of Stockholm, Sweden (EPN Dnr 2012/111-31/1; 2012/2091-32). Fresh frozen tissue was obtained through the Netherlands Brain Bank (NBB). The work with human tissues was carried out according to the Code of Ethics of the World Medical Association (Declaration of Helsinki).

### Tissue sectioning and laser capture

Sample preparation prior LCM-seq was carried out as follows. Frozen midbrain tissues obtained from the brain banks were attached to chucks using pre-cooled OCT embedding medium (Histolab). 10 μm-thick coronal sections were acquired in a cryostat at −20 °C and placed onto precooled-PEN membrane glass slides (Zeiss). For RNAscope experiments, sections were cut at 12 μm-thickness and attached to Superfrost® Plus slides (Thermo Scientific). The slides with sections were kept at −20 °C during the sectioning and subsequently stored at −80 °C until further processed.

The laser capture procedure followed by sequencing library preparation (LCM-seq) was carried out as described (Nichterwitz et al. 2016; Nichterwitz et al. 2018).

### Mapping and gene expression quantification

Samples were sequenced using an Illumina HiSeq2000 or HiSeq2500 platform (reads of 43 or 50 bp in length respectively). The uniquely mapped reads were obtained by mapping to the human reference genome hg38/GRCh38 using STAR with default settings. The reads per kilobase of transcript per million mapped reads (RPKM) were estimated using “rpkmforgenes” (Ramskold et al. 2009). As part of the quality control we verified that samples had >1 million reads and >7000 genes expressed with RPKM>1. As all samples processed exceeded this cutoff and had between 1.3∼9.8 million reads and 7.9 ∼ 12.3 kilo genes expressed with RPKM>1, all samples were included. The correlation coefficient between any two nearest samples was considered to be above 0.7. For cases having more than one replicate per group, corresponding samples were averaged before analysis so that each case had only one SNc and one VTA. Additionally, we confirmed the expression of known midbrain dopamine neuron markers and the purity of each sample (Fig. 1A).

### Differential expression analyses

Differentially expressed genes were identified using the R package “DESeq2” (version: 1.16.1) (Love et al. 2014) where the cutoff for significance was an adjusted P value of 0.05. Identified DEGs (from different analysis and summarized below) are shown in Supplemental Tables S4 and S6 to S9.

Supplemental Table S4: Mouse differentially expressed genes (La Manno et al. 2016): 390 DEGs calculated from 73 SNc and 170 VTA cells.

Supplemental Table S6: Human differentially expressed genes (current study): 74 DEGs calculated from 16 SNc and 14 VTA samples from 18 male individuals.

Supplemental Table S7: Human differentially expressed genes (Nichterwitz et al. 2016): 100 DEGs calculated from 4 SNc and 3 VTA samples from 3 female subjects.

Supplemental Table S8: Human stable genes: 33 DEGs calculated from 12 individuals (from the current study) using the bootstrapping approach.

Supplemental Table S9: Mouse stable genes: 89 DEGs calculated from 73 SNc and VTA cells (La Manno et al. 2016) using the bootstrapping approach.

### Bootstrapping approach coupled with DESeq2

To counteract the variability among human subjects and identify the most reliable DEGs between SNc and VTA neurons across datasets we developed a bootstrapping approach coupled with DESeq2 (Fig. 2A; Supplemental Fig. S4C). The stable genes output of this analysis is correlated with the sample size and give an unbiased estimation of the number of individuals required to consistently distinguish these closely related subpopulations. Importantly this computational tool can be used for the comparison of any other two groups (https://github.com/shanglicheng/BootstrappingWithoutReplacement).

#### In detail

1. Define ‥N‥ and ‥M‥ as the number of samples in Group 1 and Group 2, respectively. Choose ‥I‥ as a reference representing a given number of samples from ‥N‥ and ‥M‥.
2. Define ‥i ‥ as the number of randomly selected samples from Group1 and Group2, where i ∈{3, 4, 5, …, ‥I-1‥}. In the human dataset, as we have 12 paired samples, the i ∈{3, 4, 5, 6, 7, 8, 9, 10, 11}.
3. Pool ‥i‥ samples (temporary considered a ‥new data set‥) and calculate DEGs with DESeq2.
4. Repeat steps 2) and 3) for ‥j‥ times (set to 1000 times in this study).
5. For every round of random selection and DESeq2, save the full list of DEGs, compute and rank their frequency.
6. Set a threshold (30% ratio in this study) to consider DEGs with higher frequency as stable genes. Reliable genes appear when frequencies are above: Total times of random pooling x ratio (300 in this study). A stringent, but fair ratio can be defined by comparing the percentage of identified stable genes overlapping with the top (most significant) 10%, 20%, 30%, …, DEGs identified by DESeq2 alone.

### Bootstrapping approach applied to human samples

To reliable identify DEGs between human SNc and VTA samples, while minimizing subject variability, we selected 12 individuals (66% of the dataset, 12 out of 18 individuals) where both neuronal populations were available and sequenced. Hence, the number of randomly selected samples (‥n‥ and ‥m‥ from ‥i‥ individuals) was from three to 11 and the algorithm repeated 1000 times (Fig. 2A).

### Bootstrapping approach applied to mouse single cells

For this adult mouse dataset (La Mano et. al, 2016) we defined the groups SNc (N=73 cells) and VTA (M=170 cells comprising VTA1, VTA2, VTA3 and VTA4). To compensate the unbalance in cell number and adjust dataset representation compared to the human analysis (66%), we first randomly collected a subset of 73 VTA cells, pairing both SNc and VTA. Similarly, the number of randomly selected samples was 20, 25, 30, …, 70 and the approach repeated again 1000 times (Supplemental Fig. S4C).

### Data visualization

Data visualization was achieved using Principal Component Analysis (PCA) and Hierarchical Clustering (H-cluster). PCA was calculated with the function “prcomp” in R with default parameters. Then samples are projected onto the first two dimensions, PC1 and PC2. For H-cluster we used the R function “pheatmap” (version 1.0.12) with the clustering method of “ward.D2”.

### Accuracy calculation

To measure the power of stable genes in resolving SNc and VTA samples we calculated the clustering accuracy. This is a quantitative parameter to evaluate correct group prediction using hierarchical clustering on the first two dimensions (PC1 and PC2) of the PCA for dimensional reduction. Hierarchical clustering is performed using the ward.D2 method on the selected principal components. Ward criterion is used in the hierarchical clustering because it is based on the multidimensional variance like principal component analysis. The partitioning of samples is based on the first branch of the hierarchical tree. The genes used for PCA are different according to the evaluation purpose.

As we have two predicted groups (SNc and VTA), the minimum value for the accuracy is 50%. In details:

1. Generate PCA of SNc and VTA.
2. Hierarchically cluster the samples by using the first two principle components, PC1 and PC2. Here we can obtain a cluster tree.
3. Group the samples into predicted SNc and predicted VTA by the two main branches based on the cluster tree.
4. Define ‥N‥ and ‥M‥ as the total number of samples in Group 1 and Group 2, respectively.
5. Define ‥n‥ and ‥m‥ as the number of correctly predicted samples for Group1 and Group2, respectively.
6. Accuracy, expressed as percent, is calculated according to the formula:

Accuracy (%)= ((‥n‥+‥m‥)/(‥N‥+‥M‥)) * 100

We calculated the accuracy for six different conditions (stable gene sets) as follows:

a. Mouse stable genes applied to mouse (Fig. 4A) or human (Fig. 4G) samples.
b. Mouse stable genes excluding (*Zcchc12, Cdh13, Serpine2*) applied to mouse (Fig. 4B) or human (Fig. 4H) samples.
c. Human stable genes applied to mouse (Fig. 4C) or human (Fig. 4E) samples.
d. Human stable genes excluding (*ZCCHC12, CDH13, SERPINE2*) applied to mouse (Fig. 4D) or human (Fig. 4F) samples.
e. Human stable genes excluding three random markers applied to mouse samples (Fig. 4K).
f. Mouse stable genes excluding three random markers applied to human samples (Fig-4L).

To identify additional (but not common) stable genes from mouse or human with predicted power to the other species, we performed the above analysis e) and f). Here we ranked the mean accuracy for a given stable gene upon random exclusion with any two others.

### RNAscope staining of human tissues

RNAscope (Wang et al. 2012) was used to verify the expression of one SNc marker (*SEZ6*) and one VTA-preferential gene (*CDH13*) based on the sequencing data. In brief, midbrain sections of human fresh frozen tissue (Supplemental Table S3) were quickly thawed and fixed with fresh PFA (4% in PBS) for 1 hour at 4’C. The RNAscope 2.5 HD Assay - RED Kit (Cat. 322360) was used using manufacturer recommendations.

To evaluate the procedure in the midbrain tissue (Supplemental Fig. S3D), we first tested a negative control probe against a bacterial gene (Cat. 310043, *dapB*-C1) and a positive control probe against tyrosine hydroxylase (Cat. 441651, *TH*-C1). Once we set up the assay, midbrain sections were stained with *SEZ6* (Cat. 411351-C1) or *CDH13* probes (Cat. 470011-C1). Slides were counterstained with fresh 50% Gill Solution (Cat. GSH132-1L, Sigma-Aldrich) for 2 minutes, washed in water and dried for 15min at 60°C before mounting with Pertex (Cat. 00811, Histolab).

For every sample (n=5), we imaged 5-6 random fields within the SNc and VTA regions. On average 194.25±43.02 cells were imaged per region and staining. Pictures were made at 40X magnification using the bright-field of a Leica microscope (DM6000/CTR6500 and DFC310 FX camera). After randomization and coding of all the images, the number of dots within melanized cells (dopamine neurons) were counted using ImageJ (version 1.48) and later the average number of dots per cells determined for each region. Investigators performing the quantification were blinded to the sample, target region (SNc and VTA) and probe staining.

### Statistical analysis

For this study, statistical analyses were performed using ‥R‥. For the RNAscope analysis a paired t test (Prism 6, Version 6.0f) was used to compare the mean average dots per cell (for *SEZ6* or *CDH13* staining) between the SNc and VTA. Where applicable, individual statistical tests are detailed in the figure legends where significance is marked by P < 0.05. The number of subjects/cells used for each experiment is listed in the figure or figure legends. Results are expressed as mean ± SD or SEM as specified in the figure legend.

## DATA ACCESS

All raw and processed sequencing data generated in this study have been submitted to the NCBI Gene Expression Omnibus (GEO; https://www.ncbi.nlm.nih.gov/geo/) under accession number GSE114918. Human samples re-analyzed from the Nichterwitz study (Nichterwitz et al. 2016) are under accession number GSE76514 and mouse adult single cells (La Manno et al. 2016) can be accessed at GSE76381. Previous human microarray studies used to verify SNc stable gene expression were accessed from the European repository ArrayExpress (E-MEXP-1416) (Cantuti-Castelvetri et al. 2007) or raw data requested to Dr. Kai C. Sonntag (Simunovic et al. 2009).

A processed table with RPKMs values for the full dataset generated in this study can also be found at Mendeley Data under DOI 10.17632/b7nh33pdmg.1

## ACKNOWLEDGMENTS

We thank Professor Abdel El Manira for critical reading of and insightful comments on the manuscript. We thank all Hedlund laboratory members for fruitful discussions. Human *post mortem* tissues were kindly received from the Netherlands Brain Bank (NBB). This work was funded by grants to E.H. from Parkinsonfonden (795/15; 910/16; 991/17; 1095/18; 1192/19); EU Joint Programme for Neurodegenerative Disease (JPND) (529-2014-7500); Swedish Medical Research Council (Vetenskapsrådet) (2016-02112); and NEURO Sweden; by grants to Q.D. from Swedish Research Council (Vetenskapsrådet) (2014-2870); Svenska Sällskapet för Medicinsk Forskning and Jeanssons Stiftelser; Åke Wiberg, Karolinska Institutets Forskningsstiftelser and Stiftelsen; by grants to J.A. from Karolinska Institutets Forskningsstiftelser, Stiftelsen för ålderssjukdomar (2014-2018) and Åhlén-stiftelsen (mA5 h18; Ärende Nr. 193031 h19). J.A. was supported by a postdoctoral fellowship from the Swedish Society for Medical Research and N.K. by a postdoctoral fellowship from Hjärnfonden, Sweden.

## Author contributions

Conceptualization, E.H., Q.D. and J.A.; Methodology and Investigation, J.A., M.C., S.C., N.K., Q.D. and E.H.; Software, Formal Analysis and Visualization, S.C., J.A., and N.K.; Writing-Original Draft, J.A., S.C., Q.D. and E.H.; Writing-review and Editing, J.A., S.C., N.K., M.C., Q.D. and E.H.; Supervision and Project Administration, E.H. and Q.D.; Funding Acquisition, E.H., Q.D. and J.A.

## REFERENCES

Ambalavanan A, Girard SL, Ahn K, Zhou S, Dionne-Laporte A, Spiegelman D, Bourassa CV, Gauthier J, Hamdan FF, Xiong L et al. 2016. De novo variants in sporadic cases of childhood onset schizophrenia. Eur J Hum Genet 24: 944–948.

Barker RA, Parmar M, Studer L, Takahashi J. 2017. Human Trials of Stem Cell-Derived Dopamine Neurons for Parkinson’s Disease: Dawn of a New Era. Cell Stem Cell 21: 569–573.

Bifsha P, Yang J, Fisher RA, Drouin J. 2014. Rgs6 is required for adult maintenance of dopaminergic neurons in the ventral substantia nigra. PLoS Genet 10: e1004863.

Bossers K, Meerhoff G, Balesar R, van Dongen JW, Kruse CG, Swaab DF, Verhaagen J. 2009. Analysis of gene expression in Parkinson’s disease: possible involvement of neurotrophic support and axon guidance in dopaminergic cell death. Brain Pathol 19: 91–107.

Cantuti-Castelvetri I, Keller-McGandy C, Bouzou B, Asteris G, Clark TW, Frosch MP, Standaert DG. 2007. Effects of gender on nigral gene expression and parkinson disease. Neurobiol Dis 26: 606–614.

Cho G, Bhat SS, Gao J, Collins JS, Rogers RC, Simensen RJ, Schwartz CE, Golden JA, Srivastava AK. 2008a. Evidence that SIZN1 is a candidate X-linked mental retardation gene. Am J Med Genet A 146A: 2644–2650.

Cho G, Lim Y, Zand D, Golden JA. 2008b. Sizn1 is a novel protein that functions as a transcriptional coactivator of bone morphogenic protein signaling. Mol Cell Biol 28: 1565–1572.

Choi BH, Kim RC, Vaughan PJ, Lau A, Van Nostrand WE, Cotman CW, Cunningham DD. 1995. Decreases in protease nexins in Alzheimer’s disease brain. Neurobiol Aging 16: 557–562.

Chung CY, Seo H, Sonntag KC, Brooks A, Lin L, Isacson O. 2005. Cell type-specific gene expression of midbrain dopaminergic neurons reveals molecules involved in their vulnerability and protection. Hum Mol Genet 14: 1709–1725.

Dahlstroem A, Fuxe K. 1964. Evidence for the Existence of Monoamine-Containing Neurons in the Central Nervous System. I. Demonstration of Monoamines in the Cell Bodies of Brain Stem Neurons. Acta Physiol Scand Suppl: SUPPL 232:231–255.

Damier P, Hirsch EC, Agid Y, Graybiel AM. 1999a. The substantia nigra of the human brain. I. Nigrosomes and the nigral matrix, a compartmental organization based on calbindin D(28K) immunohistochemistry. Brain 122 (Pt 8): 1421–1436.

Damier P, Hirsch EC, Agid Y, Graybiel AM. 1999b. The substantia nigra of the human brain. II. Patterns of loss of dopamine-containing neurons in Parkinson’s disease. Brain 122 (Pt 8): 1437–1448.

Di Salvio M, Di Giovannantonio LG, Omodei D, Acampora D, Simeone A. 2010. Otx2 expression is restricted to dopaminergic neurons of the ventral tegmental area in the adult brain. Int J Dev Biol 54: 939–945.

Forero A, Rivero O, Waldchen S, Ku HP, Kiser DP, Gartner Y, Pennington LS, Waider J, Gaspar P, Jansch C et al. 2017. Cadherin-13 Deficiency Increases Dorsal Raphe 5-HT Neuron Density and Prefrontal Cortex Innervation in the Mouse Brain. Front Cell Neurosci 11: 307.

Ganat YM, Calder EL, Kriks S, Nelander J, Tu EY, Jia F, Battista D, Harrison N, Parmar M, Tomishima MJ et al. 2012. Identification of embryonic stem cell-derived midbrain dopaminergic neurons for engraftment. J Clin Invest 122: 2928–2939.

Greene JG, Dingledine R, Greenamyre JT. 2005. Gene expression profiling of rat midbrain dopamine neurons: implications for selective vulnerability in parkinsonism. Neurobiol Dis 18: 19–31.

Grimm J, Mueller A, Hefti F, Rosenthal A. 2004. Molecular basis for catecholaminergic neuron diversity. Proc Natl Acad Sci U S A 101: 13891–13896.

Hallett PJ, Cooper O, Sadi D, Robertson H, Mendez I, Isacson O. 2014. Long-term health of dopaminergic neuron transplants in Parkinson’s disease patients. Cell Rep 7: 1755–1761.

Haque NS, LeBlanc CJ, Isacson O. 1997. Differential dissection of the rat E16 ventral mesencephalon and survival and reinnervation of the 6-OHDA-lesioned striatum by a subset of aldehyde dehydrogenase-positive TH neurons. Cell Transplant 6: 239–248.

Hedlund E, Perlmann T. 2009. Neuronal cell replacement in Parkinson’s disease. J Intern Med 266: 358–371.

Hook PW, McClymont SA, Cannon GH, Law WD, Morton AJ, Goff LA, McCallion AS. 2018. Single-Cell RNA-Seq of Mouse Dopaminergic Neurons Informs Candidate Gene Selection for Sporadic Parkinson Disease. Am J Hum Genet 102: 427–446.

Kefalopoulou Z, Politis M, Piccini P, Mencacci N, Bhatia K, Jahanshahi M, Widner H, Rehncrona S, Brundin P, Bjorklund A et al. 2014. Long-term clinical outcome of fetal cell transplantation for Parkinson disease: two case reports. JAMA Neurol 71: 83–87.

Khoonsari PE, Haggmark A, Lonnberg M, Mikus M, Kilander L, Lannfelt L, Bergquist J, Ingelsson M, Nilsson P, Kultima K et al. 2016. Analysis of the Cerebrospinal Fluid Proteome in Alzheimer’s Disease. PLoS One 11: e0150672.

Kim T, Loh YP. 2006. Protease nexin-1 promotes secretory granule biogenesis by preventing granule protein degradation. Mol Biol Cell 17: 789–798.

Kirkeby A, Nolbrant S, Tiklova K, Heuer A, Kee N, Cardoso T, Ottosson DR, Lelos MJ, Rifes P, Dunnett SB et al. 2017. Predictive Markers Guide Differentiation to Improve Graft Outcome in Clinical Translation of hESC-Based Therapy for Parkinson’s Disease. Cell Stem Cell 20: 135–148.

Kriks S, Shim JW, Piao J, Ganat YM, Wakeman DR, Xie Z, Carrillo-Reid L, Auyeung G, Antonacci C, Buch A et al. 2011. Dopamine neurons derived from human ES cells efficiently engraft in animal models of Parkinson’s disease. Nature 480: 547–551.

La Manno G, Gyllborg D, Codeluppi S, Nishimura K, Salto C, Zeisel A, Borm LE, Stott SRW, Toledo EM, Villaescusa JC et al. 2016. Molecular Diversity of Midbrain Development in Mouse, Human, and Stem Cells. Cell 167: 566–580 e519.

Love MI, Huber W, Anders S. 2014. Moderated estimation of fold change and dispersion for RNA-seq data with DESeq2. Genome Biol 15: 550.

Mele M, Ferreira PG, Reverter F, DeLuca DS, Monlong J, Sammeth M, Young TR, Goldmann JM, Pervouchine DD, Sullivan TJ et al. 2015. Human genomics. The human transcriptome across tissues and individuals. Science 348: 660–665.

Miller JD, Ganat YM, Kishinevsky S, Bowman RL, Liu B, Tu EY, Mandal PK, Vera E, Shim JW, Kriks S et al. 2013. Human iPSC-based modeling of late-onset disease via progerin-induced aging. Cell Stem Cell 13: 691–705.

Mulley JC, Iona X, Hodgson B, Heron SE, Berkovic SF, Scheffer IE, Dibbens LM. 2011. The Role of Seizure-Related SEZ6 as a Susceptibility Gene in Febrile Seizures. Neurol Res Int 2011: 917565.

Nichterwitz S, Benitez JA, Hoogstraaten R, Deng Q, Hedlund E. 2018. LCM-Seq: A Method for Spatial Transcriptomic Profiling Using Laser Capture Microdissection Coupled with PolyA-Based RNA Sequencing. Methods Mol Biol 1649: 95–110.

Nichterwitz S, Chen G, Aguila Benitez J, Yilmaz M, Storvall H, Cao M, Sandberg R, Deng Q, Hedlund E. 2016. Laser capture microscopy coupled with Smart-seq2 for precise spatial transcriptomic profiling. Nat Commun 7: 12139.

Nolbrant S, Heuer A, Parmar M, Kirkeby A. 2017. Generation of high-purity human ventral midbrain dopaminergic progenitors for in vitro maturation and intracerebral transplantation. Nat Protoc 12: 1962–1979.

Panman L, Papathanou M, Laguna A, Oosterveen T, Volakakis N, Acampora D, Kurtsdotter I, Yoshitake T, Kehr J, Joodmardi E et al. 2014. Sox6 and Otx2 control the specification of substantia nigra and ventral tegmental area dopamine neurons. Cell Rep 8: 1018–1025.

Paracchini L, Beltrame L, Boeri L, Fusco F, Caffarra P, Marchini S, Albani D, Forloni G. 2018. Exome sequencing in an Italian family with Alzheimer’s disease points to a role for seizure-related gene 6 (SEZ6) rare variant R615H. Alzheimers Res Ther 10: 106.

Poulin JF, Zou J, Drouin-Ouellet J, Kim KY, Cicchetti F, Awatramani RB. 2014. Defining midbrain dopaminergic neuron diversity by single-cell gene expression profiling. Cell Rep 9: 930–943.

Ramskold D, Wang ET, Burge CB, Sandberg R. 2009. An abundance of ubiquitously expressed genes revealed by tissue transcriptome sequence data. PLoS Comput Biol 5: e1000598.

Redies C, Hertel N, Hubner CA. 2012. Cadherins and neuropsychiatric disorders. Brain Res 1470: 130–144.

Reyes S, Fu Y, Double K, Thompson L, Kirik D, Paxinos G, Halliday GM. 2012. GIRK2 expression in dopamine neurons of the substantia nigra and ventral tegmental area. J Comp Neurol 520: 2591–2607.

Schultzberg M, Dunnett SB, Bjorklund A, Stenevi U, Hokfelt T, Dockray GJ, Goldstein M. 1984. Dopamine and cholecystokinin immunoreactive neurons in mesencephalic grafts reinnervating the neostriatum: evidence for selective growth regulation. Neuroscience 12: 17–32.

Simunovic F, Yi M, Wang Y, Macey L, Brown LT, Krichevsky AM, Andersen SL, Stephens RM, Benes FM, Sonntag KC. 2009. Gene expression profiling of substantia nigra dopamine neurons: further insights into Parkinson’s disease pathology. Brain 132: 1795–1809.

Sonntag KC, Song B, Lee N, Jung JH, Cha Y, Leblanc P, Neff C, Kong SW, Carter BS, Schweitzer J et al. 2018. Pluripotent stem cell-based therapy for Parkinson’s disease: Current status and future prospects. Prog Neurobiol 168: 1–20.

Studer L. 2017. Strategies for bringing stem cell-derived dopamine neurons to the clinic-The NYSTEM trial. Prog Brain Res 230: 191–212.

Thompson L, Barraud P, Andersson E, Kirik D, Bjorklund A. 2005. Identification of dopaminergic neurons of nigral and ventral tegmental area subtypes in grafts of fetal ventral mesencephalon based on cell morphology, protein expression, and efferent projections. J Neurosci 25: 6467–6477.

Vera E, Bosco N, Studer L. 2016. Generating Late-Onset Human iPSC-Based Disease Models by Inducing Neuronal Age-Related Phenotypes through Telomerase Manipulation. Cell Rep 17: 1184–1192.

Wang F, Flanagan J, Su N, Wang LC, Bui S, Nielson A, Wu X, Vo HT, Ma XJ, Luo Y. 2012. RNAscope: a novel in situ RNA analysis platform for formalin-fixed, paraffin-embedded tissues. J Mol Diagn 14: 22–29.

Yu ZL, Jiang JM, Wu DH, Xie HJ, Jiang JJ, Zhou L, Peng L, Bao GS. 2007. Febrile seizures are associated with mutation of seizure-related (SEZ) 6, a brain-specific gene. J Neurosci Res 85: 166–172.

Zhang Y, Chen K, Sloan SA, Bennett ML, Scholze AR, O’Keeffe S, Phatnani HP, Guarnieri P, Caneda C, Ruderisch N et al. 2014. An RNA-sequencing transcriptome and splicing database of glia, neurons, and vascular cells of the cerebral cortex. J Neurosci 34: 11929–11947.

